# PvdL Orchestrates the Assembly of the Nonribosomal Peptide Synthetases Involved in Pyoverdine Biosynthesis in *Pseudomonas aeruginosa*

**DOI:** 10.1101/2023.12.12.571276

**Authors:** Hanna Manko, Tania Steffan, Véronique Gasser, Yves Mély, Isabelle J Schalk, Julien Godet

**Affiliations:** Laboratoire de BioImagerie et Pathologies, UMR CNRS 7021, ITI InnoVec, Université de Strasbourg, Illkirch, France; CNRS, UMR 7242, ITI InnoVec, ESBS, Illkirch, Strasbourg, France; Laboratoire de BioImagerie et Pathologies, UMR CNRS 7021, Université de Strasbourg, Illkirch, France; Faculté de Pharmacie, Université de Strasbourg, Illkirch, France; Groupe Méthodes Recherche Clinique, Hôpitaux Universitaires de Strasbourg, France; Laboratoire iCube, UMR CNRS 7357, Equipe IMAGeS, Université de Strasbourg, Strasbourg, France

**Keywords:** Pseudomonas aeruginosa, NRPSs, pyoverdine, super-resolution microscopy, DNA-PAINT, colocalization, sptPALM, FLIM-FRET

## Abstract

The pyoverdine siderophore is produced by *Pseudomonas aeruginosa* to access iron. Its synthesis involves the complex co-ordination of four nonribosomal peptide synthetases (NRPSs), responsible for assembling the pyoverdine peptide backbone. The precise cellular organization of these NRPSs and their mechanisms of interaction remain unclear.

Here, we used a combination of several single-molecule microscopy techniques to to elucidate the spatial arrangement of NRPSs within pyoverdine-producing cells. Our findings revealed that PvdL differs from the three other NRPS in term of localization and mobility patterns. PvdL is predominantly located at the inner membrane, while the other also explore the cytoplasmic compartment. Leveraging the power of multicolor single-molecule localization, we further reveal co-localization between PvdL and the other NRPSs, suggesting a pivotal role for PvdL in orchestrating the intricate biosynthetic pathway.

Our observations strongly indicates that PvdL serves as a central orchestrator in the assembly of NRPSs involved in pyoverdine biosynthesis, assuming a critical regulatory function.

## Introduction

Pseudomonas species exhibit remarkable adaptability to diverse environments associated with human activities due to their versatile metabolic capabilities (1). Specific metabolic pathways can be activated in response to environmental conditions, enabling bacteria to generate a multitude of distinct secondary metabolites that confer fitness or selective advantages (2). Non-ribosomal peptides (NRPs) are a large and diverse family of secondary metabolites that often serve specialized functions such as defence, communication, or colonization (3, 4). NRPs are synthetized by Non-Ribosomal Peptide Synthetases (NRPSs), a family of high molecular-weight modular enzymes that can produce peptide molecules independently from ribosomes (5, 6). While ribosomal peptide synthesis is limited to 20 natural amino acids, NRPSs have the ability to use a diverse array of building blocks (7), enabling the synthesis of an extensive spectrum of secondary metabolites, encompassing toxins, virulence factors, and molecules with therapeutic interests (6). In *P. aeruginosa*, the peptide backbone of pyoverdine is synthesized by four distinct NRPSs. Pyoverdine is the main siderophore produced by *P. aeruginosa* (8). Siderophores are structurally diverse compounds with molecular weight between 200 and 2000 Da, produced by pathogenic and nonpathogenic bacteria and fungi in order to get access to iron, a nutrient essential for cellular growth (9). Owing to their capability to immobilize iron, siderophores can effectively vampirize iron to deprive competing bacteria or the host of this essential nutrient. (10–13). Blocking the production of pyoverdine or using inhibitors against as been reported as sufficient to mitigate *P. aeruginosa* pathogenesis (14, 15). Siderophores are also important mediators of interactions between members of microbial assemblies leading to cooperative, exploitative and competitive interactions between individuals (16). Pyoverdine is an example of mediator for local mutualistic cooperation providing direct benefits to producers carrying the cost of production but also to non-producing but recipient cells (17). Siderophores can also act as important mediators of interactions with eukaryotic hosts (18).

The pyoverdine biosynthesis process starts in the cytoplasm with the synthesis of a precursor of pyoverdine further maturated in the bacterial periplasm before secretion (Figure 1). In the cytoplasm the synthesis involves a relatively intricate enzymatic system comprising four distinct NRPSs: PvdL, PvdI, PvdJ and PvdD and three enzymes (PvdH, PvdA and PvdF) producing modified amino acids. PvdL catalyzes the attachment of a fatty acid to a glutamate residue, followed by the addition of L-tyrosine and L-Dab to the product. The resulting 3 amino-acid product is transferred to PvdI, which adds L-Serine, L-Arginine, L-Serine and L-hydroxyornithine, and to PvdJ which adds L-Lysine and L-Hydroxyornithine. Finally, PvdD adds of two more L-Threonines and cyclizes the pyoverdine backbone (8, 19). Several attempts were done to reveal the cellular organization of the enzymes involved in the pyoverdine biosynthesis pathway, leading to the suggestion that NRPSs may form membrane-bound multi-enzymatic complex, called the siderosomes (20–22). Enzymes responsible for the biosynthesis of modified amino-acids would also be part of the siderosomes. However, a clear bio-chemical characterization of siderosomes is missing. So far, they have not been isolated nor reconstituted *in vitro*, possibly due to the large molecular weight of the NRPSs making them challenging to express and purify or simply because siderosomes form transiently in cells. These reasons might limit the investigation of siderosomes in their native environment. *In cellulo* FRET-FLIM and single molecule tracking revealed, for example, that the ornithine hydroxylase PvdA physically interacts with all four NRPS, albeit with different stoichiometries depending on whether or not the NRPSs use the modified amino acids produced by PvdA (23).

**Fig. 1.**
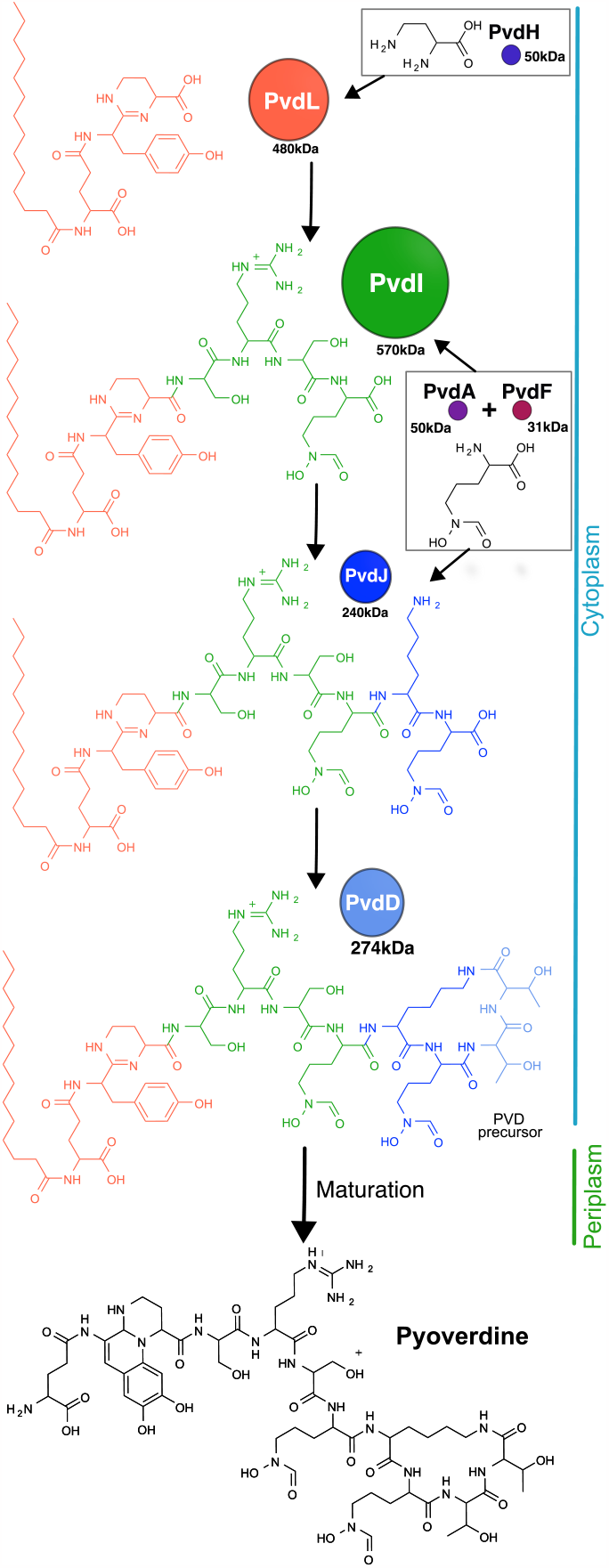
Assembly of the pyoverdine molecule. The cytoplasmic pyoverdine precursor is assembled by four NRPSs (PvdL, PvdI, PvdJ and PvdD) with three smaller enzymes (PvdH, PvdA and PvdF) providing modified amino-acids. Each NRPS adds blocks (like amino-acids) in the peptide backbone. At the very first step, PvdL introduces myristic acid which is removed once the pyoverdine precursor is transported into the periplasm and before it undergoes maturation of the chromophore.

Elaborate biosynthesis pathways, characterized by multiple sequential steps, have to be accurately regulated in order to effectively balance the competing imperatives of minimizing production costs while simultaneously optimizing reaction yields (8). The regulation of these complexes is likely to involve the timely expression levels of the different enzymes, as well as their temporal stabilities and spatial organization. However, the spatial organization and dynamics of these four NRPSs in cells have received only minimal attention in the literature. More generally, data focusing on the spatial and temporal regulation of biosynthetic pathways in their native environment is scarce (24).

In this work we proposed to use a combination of single-molecule microscopy techniques to gain insight into the intracellular organization in cells of the four NRPSs of the pyoverdine biosynthesis pathway. We showed that PvdL has a localization and mobility pattern distinct from the other three NRPSs. PvdL is mainly localized at the inner membrane while the others also explore the cytoplasmic compartment, thus having two diffusions regimes, one of which is very similar to that of PvdL. Leveraging the power of multicolor single-molecule localization microscopy, we further reveal co-localization between PvdL and the remaining NRPS. The unique localization and mobility patterns of PvdL suggest that it may play a key role in serving as an anchor point for other NRPSs during the dynamic process of pyoverdine biosynthesis.

## Results

### Localizations of NRPSs in cells

Bacteria have evolved diverse mechanisms to address challenges associated with efficient growth and replication (25, 26). Precise regulation, including spatial control, of the complex and resource-intensive metabolic pathways appears to be essential. Therefore, we sought to investigate the intracellular spatial localizations of the fours NRPSs of the pyoverdine pathway using single-molecule localization DNA-paint microscopy.

We used *Pseudomonas aeruginosa* PAO1 strains in which either PvdL, PvdI, PvdJ, or PvdD were fused to eGFP at the chromosomal level. The expression of the fluorescent enzymes was induced by iron-deficient growing conditions similarly to the wild-type PAO1 (27). In fixed cells, the eGFP moiety served as the target for immunostaining, where an anti-eGFP primary antibody was subsequently recognized by a secondary antibody containing a DNA-PAINT docking strand. In DNA-PAINT, the addition in the sample of short dye-labeled (‘imager’) oligonucleotides transiently binding to their complementary target (‘docking’) strands creates the necessary ‘blinking’ to enable stochastic single-molecule localization microscopy (28). This approach offers several advantages compared to the direct imaging of labelled secondary antibodies, including the predictability of DNA binding and unbinding events, coupled with minimal (virtually no) photo-bleaching (29). This combination facilitates accurate localization of NRPS localizations, as a single target protein is associated with multiple and repeatedly detectable fluorescence bursts within the sample.

Cell imaging was performed in three dimensions (3D) to reduce artefacts resulting from the two-dimensional projections of three-dimensional samples during 2D imaging. For the z-axis measurement, we used a cylindrical lens to introduce astigmatism into the imaging system. The z-encoded orientation of the PSF was then measured by fitting an ellipse to the image of the fluorophore. This approach allowed to selectively build ∼ 100 nm slices within the central portion of the bacteria and to compute a cross-sectional view (Figure 2). Interestingly, discrete punctate localizations along the cellular structure were observed for the four NRPSs. Cross-sectional observations revealed partial exclusion zones within the central region of the cell. This region probably corresponds to the area of the nucleoid (Figure 2, a, b, cross-section views).

**Fig. 2.**
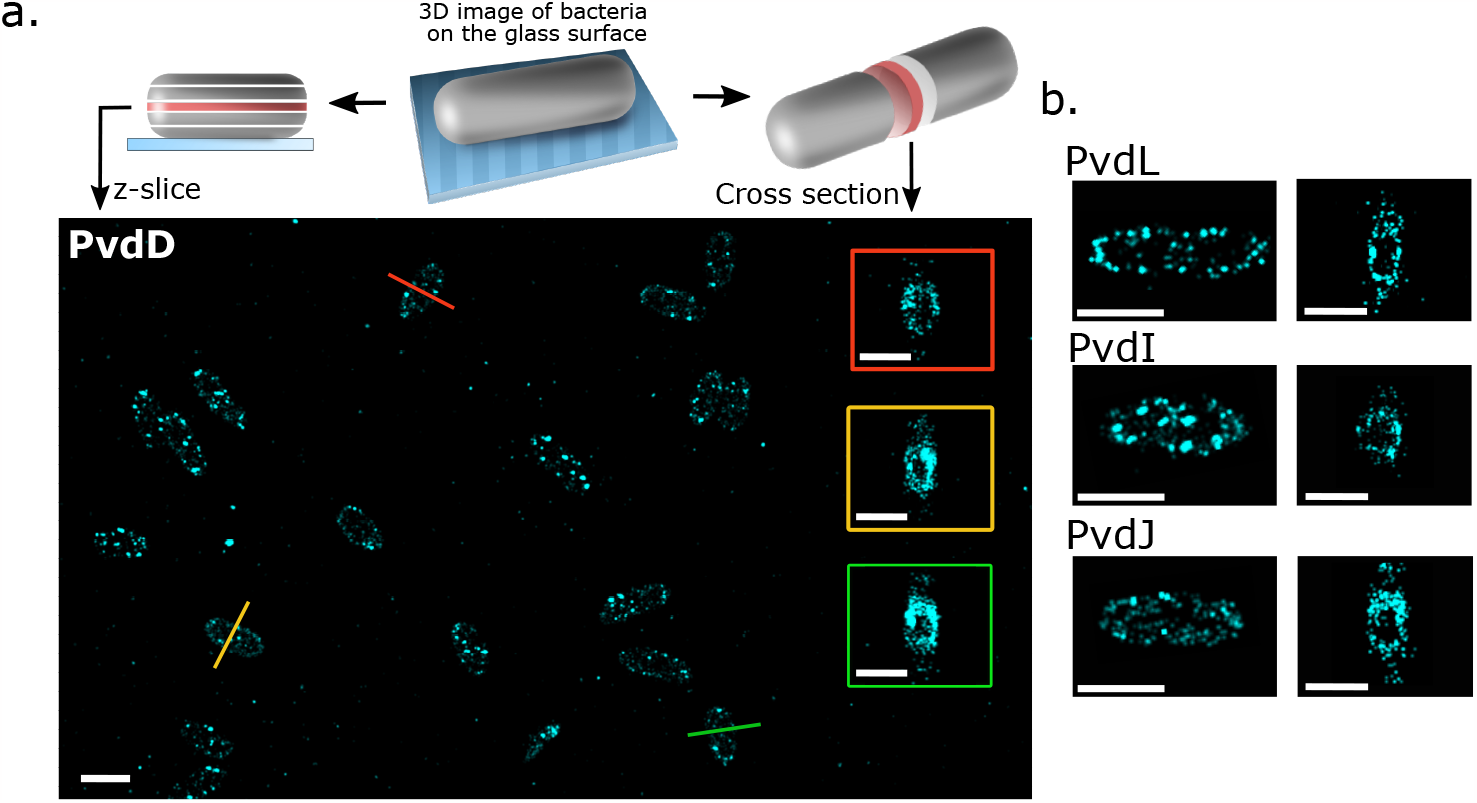
**a**. Representative z-axis slice projections of 3D localizations of labeled PvdD in fixed bacterial cells on a glass coverslip (scale bar = 2 *μ*m). Some cross sections of bacteria are presented in the colored boxes (scale bar = 1*μ*m). **b**. z-projections and cross sections obtained for PvdL, PvdI and PvdJ in fixed *P. aeruginosa* cells (scale bars = 1 *μ*m).

Previous reports showed exclusion of large proteins from the dense nucleoid region (30, 31). But a nucleoid exclusion has not been described before for proteins of the pyoverdine pathway, neither by FRAP (22) nor by 2D-sptPALM experiments (23) - probably because much smaller proteins were tracked. The discrete distribution of the NRPSs throughout the cell is an interesting finding. It can ensure that NRPSs are equitably distributed to the daughter cells during replication. It can also increase the surface area for interactions with other molecules, or create local microenvironments that are conducive to pyoverdine production.

Nevertheless, these data obtained with the DNA-PAINT technique were performed on paraformaldehyde-fixed bacteria. While we assume that neither fixation nor immunostaining perturbs the distribution of NRPSs in cells, it is important to note that it provides a static picture of the enzymes localization. To overcome this limit, we sought to examine the diffusion patterns of NRPSs in living cells.

### Diffusion of NRPSs in living cells

To explore the diffusion of NRPSs in their native environment, we generated strains expressing either PvdL, PvdI, PvdJ or PvdD fused to a photoactivatable mCherry (PAm-Cherry) at the chromosomal level. We performed single particle tracking localization microscopy (sptPALM) (32) with high oblique illumination on living cells expressing the NRPS and that were immobilized on an agarose pad. The tracking data set was derived from a minimum of four independent experimental replicates. Once the localizations were determined frame by frame with sub-diffraction precision, the molecular trajectories were reconstructed based on Linear Assignment Problem (LAP) tracker algorithm (33, 34). The trajectories were clustered by cell to create cellular diffusion maps (Figure 3 a).

**Fig. 3.**
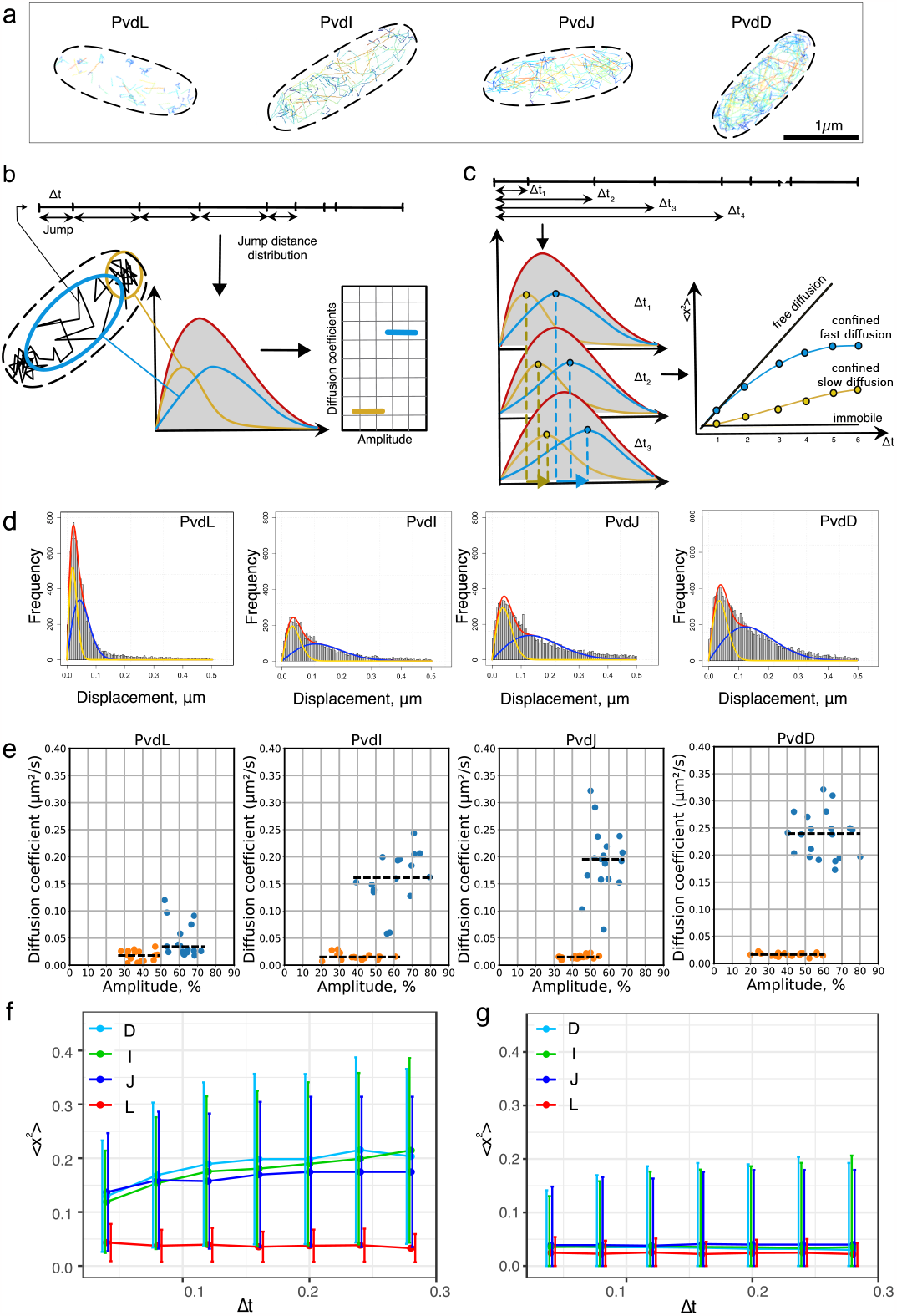
**a Diffusion maps.** Representative diffusion maps for PvdL, I, J and D illustrate differences in the number of trajectories and explored surface areas. **b Jump distance analysis principle** Jumps are computed from trajectories. The jump distributions can be used to obtain diffusion coefficients and the relative amplitude of the different diffusing subpopulations. **c Mean Square displacement** By varying the elapsed time, new jumps can be computed and the mean square displacement of the different subpopulations of diffusing molecules can be estimated. **d. Experimental jump distance histograms** corresponding to one field of view in the sample. The histograms were fitted using a two component model, with fast (blue line) and slow (orange line) diffusion regimes. **e Diffusion parameters** The diffusion coefficients and the corresponding amplitudes of the fast (blue dots) and the slow (orange dots) diffusion modes were determined. Each dot corresponds to the coefficient calculated by analysing all the tracks containing more than three spots in one field of view. The median values of the diffusion coefficient are shown as black dashed lines. **f Mean square displacement** MSD (median *±* sd) as a function of Δ*t* for the fast (left) and slow(right) regimes of diffusion of the NRPSs.

Comparison of the four representative diffusion maps (Figure 3 a) indicates that the density of the trajectories differs between the different NRPS, in accordance with their relative expression in the cells - PvdL being the least expressed and PvdD being the most expressed. This visual impression was further reinforced by the fact that the areas explored by the trajectories were not the same. As represented by the jump size color code, PvdL diffusion was the most restricted (small jumps) with low-exploration traces remaining close to the cell boundary. On the contrary, PvdI, PvdJ and PvdD showed different regimes of diffusion, with low-exploration traces at the outline of the cell coexisting with traces exploring much more surface inside the cell (Figure 3 a). To provide further quantitative insight, we analyzed these data using jump distance analysis. Jump distance analysis calculates the distances between the positions of molecules in consecutive frames, each spanning a delta time corresponding to the time elapsed between the two frames (Δ*t* =40 ms) (35). These distances are then plotted into a histogram to estimate a jump distance distribution. The jump distance distributions provide information about the diffusion coefficient of the molecules and facilitates the identification of distinct subpopulations of molecules diffusing at different rates in the sample (Figure 3 b). In addition, we extended the analysis to different delta times (3, 4, 5 frames, and so forth) to investigate whether molecules were undergoing confined diffusion and to distinguish between extremely slow diffusion and immobility (Figure 3 c). Varying the delta time allows the JD analysis to converge towards a mean square displacement (MSD) analysis, but with the ability to estimate a MSD for each subpopulation of molecules.

The best fits to describe the NRPS jump distance distributions were obtained for two-population models (Figure 3 d). Interestingly, the values of the diffusion coefficients for the slow diffusion mode were similar for all four NRPSs (see orange points in Figure 3, e.). The diffusion coefficients corresponding to the fast mode were found to be slightly different between PvdI, PvdJ and PvdD, and strongly different from that of PvdL. The fast diffusion coefficients of PvdI, L and D seemed to differ independently of their molecular weight (Figure 3 e, blue points). The diffusion mode in the bacterial cytoplasm is difficult to interpret from the values of the diffusion coefficient, but they suggest that NRPSs could diffuse as complexes or multimers. The slowest diffusion rate was found for PvdL with a diffusion coefficient of 0.05 [0.02–0.06 ] μm^2^.s^−1^ (mean, [IQR]) (Figure 3 e, blue points). The diffusion coefficients of the two diffusion modes of PvdL were found to be close (Figure 2 c), allowing the histogram to be fitted adequately with a single population in some cells. This suggests that PvdL is mostly confined to the inner membrane with rare exploration of the cytoplasmic compartment.

Even when bound to the inner membrane, the diffusion of PvdL is slow. This was confirmed by the MSD curves corresponding to the slow component, which did not increase with increasing Δt (Figure 3 g). This shows that a fraction of PvdL appeared immobile in the membrane when tracked with a pointing precision of about 30 nm. In line with the fact that the jump distribution could almost be considered a single population, the MSD of the faster component also remains constant over the Δt range (Figure 3 f). On the contrary, PvdI, J and D showed a MSD compatible with confined diffusion, as expected from the dimension of the cytoplasmic compartment in which these proteins can diffuse (36). Interestingly and like PvdL, their slow component appeared to be immobile. Despite the limitations of the 2D projections, the locations of the immobile tracks suggest that slow diffusion regimes occur mostly near the inner membrane (see supplementary Figure Sx).

Taken together, these results suggest that PvdL behaves differently from the three other NRPSs. PvdL is almost exclusively associated to the inner membrane with very rare excursions into the cytoplasm. In contrast, PvdI, J, and L diffuse in the cytoplasm. They also experienced a slow diffusion mode that is very similar to that of PvdL, suggesting that the different NRPSs assemble in large complex at the inner membrane. Therefore, we sought to investigate direct NRPS-NRPS interaction in living cells.

### NRPS-NRPS interactions

To explore direct NRPS-NRPS interactions in living cells, we used Förster resonance energy transfer measured by fluorescence lifetime imaging (FLIM-FRET) (37). FLIM-FRET allows the measurement and the mapping of the fluorescence lifetime of a donor fluorophore directly in living cells. If FRET occurs, the fluorescence lifetime of the donor fluorophore decreases due to the non-radiative energy transfer to an acceptor fluorophore that is located in close proximity (typically in the range 1 to 10 nm). The strong distance dependence of FRET (∝1/R^6^) means that it is only possible for FRET to occur when the donor and acceptor fluorophores are located very close to each other (< 10 nm). At the level of proteins, this almost always corresponds to situations where physical interactions occur. Therefore, FLIM-FRET is particularly suitable for measuring direct protein-protein interactions.

We used different combination of protein mutants in which one NRPS was fused to eGFP (donor) and another to mCherry (acceptor) (Table 1). We also used bacteria expressing the appropriate NRPS tagged with eGFP only as control (donor only). In these combinations, PvdL was systematically one of the two NRPSs that was tagged with a fluorescent protein. The lifetime values of donor alone (*τ*_*d*_) and in the presence of acceptor (*τ*_*da*_) are shown in table Table 1. The change in lifetime due to acceptor was the most pronounced for the PvdL/PvdD pair. It was associated to a clear shift in the position of the lifetime phasor (Figure 4), confirming an interaction between the two proteins. The lifetime values did not change when the acceptor was present for the PvdL/PvdJ pair. The decrease of the donor lifetime in the PvdL/PvdI was limited, with a FRET efficiency almost indistinguishable from noise. Therefore, it was not possible to evidence FRET for an NRPS pair other than PvdL/PvdD. Nevertheless, due to the large size of NRPSs, these data cannot completely rule out the presence of direct interactions between these proteins, even in the absence of FRET. Indeed, if the presence of FRET indicates interaction, the absence of FRET does not necessarily indicate an absence of interaction because the two fluorescent tags can fall too far apart in the complex. Ideally, NRPS labelling should have been performed at positions other than the N- or C-termini to increase the likelihood of measuring FRET when proteins interact. However, without structural understanding of potential interaction sites, internal labelling seemed arbitrary. There is also a risk of losing NRPS functionality. We therefore chose to investigate potential interactions between NRPSs using co-localisation measurements.

**Table 1.** The average lifetime values for donors (*τ*_*d*_), donor-acceptor pairs (*τ*_*da*_), and FRET efficiency (E) observed through FLIM-FRET experiments for PvdL and the three other NRPSs.

| Strain name | $\tau_d$ (ns) | $\tau_{da}$ (ns) | E |
| --- | --- | --- | --- |
| eGFP-PvdL / PvdJ-mCherry | 2.3 | 2.3 | 0.00 |
| PvdJ-eGFP / mCherry-PvdL | 2.3 | 2.3 | 0.00 |
| eGFP-PvdL / mCherry-PvdD | 2.3 | 2.0 | 0.13 |
| eGFP-PvdL / PvdI-mCherry | 2.3 | 2.2 | 0.04 |

**Fig. 4.**
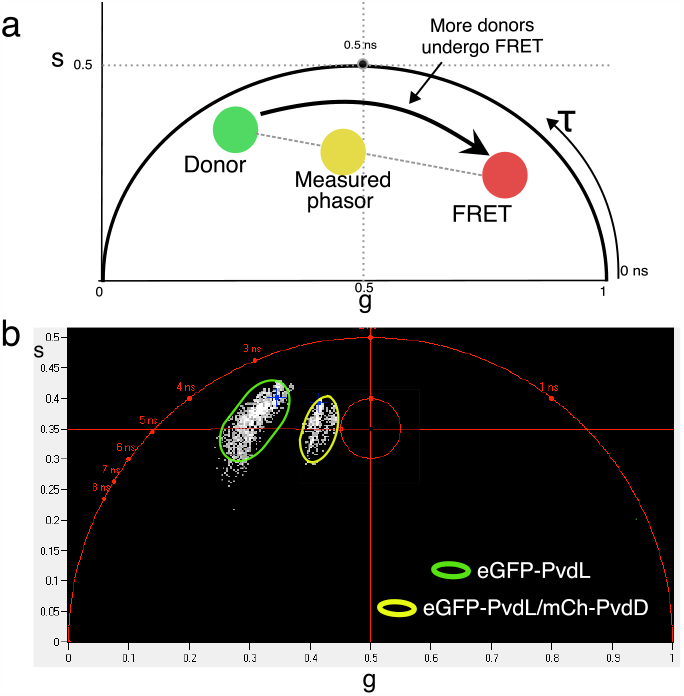
**a. Phasor approach principle.** On a phasor plot, each phasor point is obtained by transformation of the fluorescence decay of one FLIM pixel. The phasor coordinates are defined by real (g) and imaginary (s) parts of the Fourier transform. The position of the donor in the phasor plot will depend on its lifetime (green filled circle). When all the donor molecules undergo FRET with high transfer efficiency, the phasor is shifted in the direction of lower lifetimes (red filled circle). In the case of lower FRET efficiency the phasor will stand on the line joining the two previously described positions (orange filled circle). **b. Phasor plot**. A clear shift between phasors of the donor (eGFP-PvdL, green oval) and the donor acceptor pair (eGFP-Pvdl/mCh-PvdD, orange oval) can be seen, showing protein-protein interaction between PvdL and PvdD.

### Co-localizations of NRPSs

For co-localization experiments, we used the same *P. aeruginosa* double mutants as for FLIM-FRET experiments, but we fixed and permeabilized the cells to allow DNA-PAINT immunolabelling. We performed dual-color DNA-PAINT with imager strands labelled with Cy3B or Atto655, which specifically target docking strands attached to an anti-mouse secondary antibody targeting anti-eGFP antibody and to an anti-rabbit secondary antibody targeting anti-mCherry antibody, respectively. Samples were imaged for 60,000 frames with an exposure time of 80 ms per frame, using focus drift control and alternating illumination sources every 20,000 frames. Localizations were extracted from raw images. When localizations were detected over several consecutive frames, the signals were merged into a single detection event. Retrieved localizations were then plotted with false color according to their channels.

As shown in Figure 5 a, the localizations observed in both channels coincided partially. To provide a quantitative measure of the co-localization, we calculated the Mander’s overlap coefficients (MOCs) (38) directly from the localizations using an approach based on Voronoi diagrams (39). The MOC is a measure of how much two signals co-occur spatially, indicating the extent to which they are localized in the same cellular or subcellular regions. It can range from 0 to 1, where 0 indicates no overlap and 1 indicates perfect overlap (Figure 5b). Two coefficients, M1 and M2, can be calculated. They represent the proportion of one signal that overlaps with the other signal, normalized by the total intensity of one of the two signals. The dissimilarities between M1 and M2 values are informative about the asymmetry in the spread of co-localizations or may express differences in the level of expression of the proteins. Compared to pixel-by-pixel calculations, MOC calculated from localizations allows more robust determination because the position of fluorescent molecules is less likely to be affected by background noise than pixel intensity values. Mander’s overlap coefficients (39) were calculated in multiple fields of view for each double mutant. For the eGFP-PvdL / PvdJ-mCherry pair, the median MOC was about 0.30 for M1 and for M2, with interquartile ranging from 0.2 to 0.5 for M1 and 0.1 to 0.5 for M2. This indicates that a significant fraction of PvdL proteins (about one third) colocalized with PvdJ (Figure 5 c). This was intriguing because no FRET could be seen between these two proteins. To confirm this observation, the same calculations were performed for the PvdJ-eGFP/mCherry-PvdL pair. Interestingly, the median MOCs were about 0.2 and 0.25 confirming that these two proteins colocalize (Figure 5 c). In line with the direct interactions evidenced by FRET, median M1 and M2 values for PvdL/PvdD were about 0.4. It indicates that about 40% of the signal of PvdD is colocalizing with PvdD, and *vice versa* - in full line with the amplitude of the low diffusing species for both PvdL and PvdL (Figure 3 e). More surprisingly, MOC values for PvdL/PvdI were even higher with both M1 and M2 about 0.6 (Figure 5 c).

**Fig. 5.**
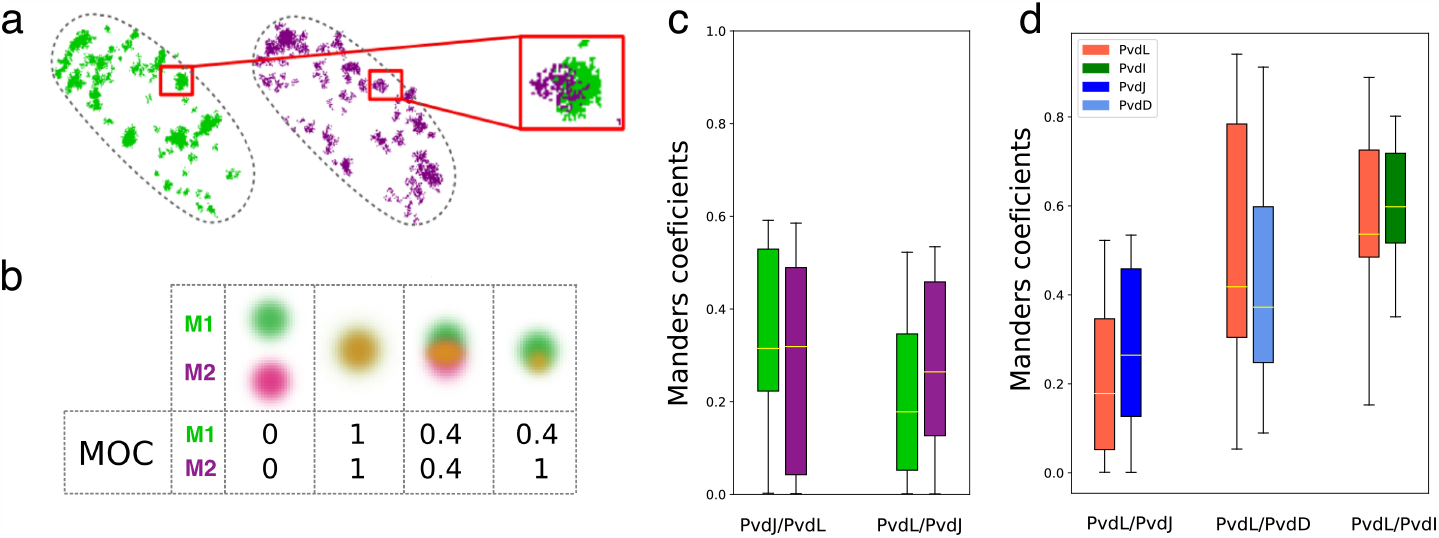
**a. Two color localization microscopy** Scatter plot of the localizations of single eGFP-PvdL (green) and PvdJ-mCherry (purple) in a cell showing partial colocalization of the two enzymes. **b. Examples of Manders overlap coefficients**. MOC range from 0 to 1. Asymetric MOC values can be observed and are informative about the asymmetry in the spread of co-localizations. **c. M1 and M2 Manders overlap coefficients calculated for eGFP-PvdL / PvdJ-mCherry and PvdJ-eGFP/mCherry-PvdL** **d. Manders overlap coefficients obtained for PvdL with the three other NRPSs**. The highest values were obtained for the PvdL/PvdD and PvdL/PvdI pairs.

Taken together, PvdL partially co-localizes with all three NRPSs. This finding is in line with the similarity observed between the diffusion coefficients of the slowly diffusing sub-populations of each NRPS and that of PvdL. This strongly suggests that PvdL could have a significant role in coordinating NRPS assembly within the pyoverdine pathway, thus assuming a central regulatory function.

## Disscussion

In this work we used a combination of several super-resolution microscopy techniques to study the organization of NRPSs in cells producing pyoverdine. We found that, in addition to the colocalization of subpopulations of PvdI, PvdJ and PvdD with PvdL, all these enzymes share a common diffusion regimen with very low mobility. We also evidenced direct PvdD-PvdL interaction in live cells. These observations suggest the formation of complexes aggregating all the NRPSs involved in biosynthesis pathway of the cytoplasmic pyoverdine precursor, and that these complexes coexist with free NRPS fractions. The free fractions of PvdI, PvdJ and PvdD mainly diffuse in the cytoplasm, whereas the free fraction of PvdL diffuses almost exclusively at the level of the inner membrane. This organization is fully compatible with the existence of transient siderosomes producing pyoverdine in cells (8, 20, 21).

Cellular reactions within metabolic pathways often occur in membrane-associated multienzyme complexes, not by free-floating enzymes (40). The diffusion rate of PvdL strongly suggests the protein is associated to the membrane. Immobilized enzymatic engineering provided evidence that spatially organized enzymatic complex achieve kinetic benefits including preventing intermediate diffusion, improving product yield, and controlling the flux of metabolites (41). The aggregation of different NRPSs by PvdL is expected to contribute to the improved efficiency and specificity of pyoverdine biosynthesis, potentially facilitating more efficient substrate channeling. The presence of a fatty acid chain introduced by PvdL at the beginning of the synthesis of pyoverdine may contribute to this process (42).

The question on how PvdL is associated to the inner membrane still needs to be addressed. Unlike some bacterial NRPSs, PvdL sequence does not contain transmembrane domains (7). PvdA, a tailoring enzyme involved in pyoverdine biosynthesis, could potentially function as an anchoring enzyme, tethering the enzymatic complex to the membrane, similar to some signaling proteins (43). PvdA possesses a hydrophobic, inner-membrane-anchoring domain at its N-terminus, facilitating its association with the inner membrane (20, 44). In addition, PvdA has been demonstrated to interact with PvdL and the three other NRPSs in live cells with multiple PvdA molecules binding to the NRPSs (23). NRPS membrane association likely serves functions such as enzyme protection, precursor coordination (45), or peptide export efficiency. In this work, we did not explore the interaction of PvdL with PvdE, an ABC transporter responsible for the export of the acylated ferribactin (the pyoverdine cytoplasmic precursor) into the periplasm (19). It would be intriguing to determine whether membrane-associated proteins or accessory proteins could not only aid PvdL membrane localization but also facilitate peptide export.

PvdL appears to play a pivotal role by cornerstoning the organization of the siderosome, not only by anchoring it to specific sites in the cell but also by regulating its behavior in term of diffusion. The advantage for a dynamic complex, with free protein fractions exchanging with the complex, over a stable complex, is not very apparent. However, advantages like sensitivity to changes in the environment to allow the cell to respond more quickly can be considered. Then, this organization might be involved in regulation, as fine control of the expression of PvdL alone might be sufficient to disrupt the efficiency of the entire process. Interestingly, the gene for PvdL is located separately from the genes for the other NRPSs (46). PvdL could then be subject to distinct regulatory mechanisms, allowing for more fine-tuned control of its expression and PvdL’s expression could be more sensitive to certain environmental cues or bacterial states. However, exploring this regulation is challenging, because delta or deficient mutants for the pyoverdine biosynthesis disrupt the positive feedback stimulating the expression of the whole pathway. Indeed, the pyoverdine metabolic pathway is expressed in conditions where iron level is low but also when ferripyoverdine is sensed by its uptake transporter FptA, leading to the release of the PvdS and PvdI sigma factors, both activating the transcription of the genes encoding the different enzymes involved in pyoverdine biosynthesis. Investigating this regulatory mechanism could deepen our understanding of how Pseudomonas bacteria selectively switch the acquisition of iron from the pyoverdine pathway to one of its multiple other iron uptake pathways in response to changes in environmental iron availability (10). This last point could be an important factor in prioritizing the expression of certain uptake pathways in the context of antibiotics which would be vectorized via iron acquisition pathways (47).

In summary, single molecule observations have revealed that PvdL plays a pivotal role in orchestrating the assembly of NRPSs involved in pyoverdine biosynthesis, suggesting a central regulatory function that will be interesting to explore.

## Materials and Methods

### Bacteria mutants, cell culture and labelling

The *Pseudomonas aeruginosa* strains used in this study are listed in supplementary materials (table S1). Mutants construction was performed as described in detail in (27). For imaging, cells were grown in lysogeny broth (LB) (L3152 Sigma Aldrich) at 30 °C under 200 rpm orbital shaking for 24 hours. The expression of the pyoverdine pathway was induced by changing the culture media for Succinate media (SM). The SM composition of this iron-deprived media was 6 g.L^−1^ K_2_HPO_4_, 3 g.L^−1^ KH_2_PO_4_, 1 g.L^−1^ [NH4]_2_SO_4_, 0.2 g.L^−1^ MgSO_4_ 7H_2_O and 4 g.L^−1^ sodium succinate, with the pH adjusted to 7.0 by adding NaOH. Cells were grown for 24 hours in SM, before being diluted 10 times and grown for an additional 24 hours in fresh SM. Before measurements, the density of cells was controlled by measuring the optical density (OD) at 600 nm. The bacteria were diluted to OD_600*nm*_ ∼ = 0.1 and grown for 2 additional hours to reach OD_600*nm*_ [0.4-0.5]. Finally cells were immobilized on a 1 % agarose pad and used for sptPALM and FLIM-FRET experiments.

For DNA-PAINT microscopy, the cells were additionally fixed with 4% formaldehyde (Sigma-Aldrich) in PBS and permeabilized with lysozyme. Targets (eGFP in single-color experiments and eGFP/mCherry in two-color) were labeled using DNA-PAINT Massive-AB-2-Plex labelling kit (Massive Photonics) according to the manufacturer’s protocol.

### Microscopy and Analysis

SptPALM and DNA-PAINT experiments were performed on a home-built bespoke Olympus IX-81 inverted optical microscope controlled by Micro-Manager software. The microscope was equipped with a 100X 1.4 NA oil-immersion objective (Olympus-Japan) and with a z-drift control and auto-focus system (ZDC Olympus). The illumination was provided by 405 nm diode laser (Oxxius), 561 nm CW diode laser (Oxxius) and 638 nm diode pumped laser (Cobolt 08-01 series) through the objective in epi-illumination or HILO illumination. The fluorescence emission was collected through the same objective and sent to the camera using a dichroic mirror (Di03-R561-t1 / Di 650-Di01 -Semrock). Before the signal was detected by 512 × 512 pixels electron multiplied charged couple device (EMCCD) camera (ImageEM-Hamamatsu Photonics - Japan) using an EM gain of 400, the fluorescence signal was filtered by longpass filter (Lp02-568RU-25 / 645LP Edge Basic, Semrock).

In sptPALM experiments the sample was simultaneoustly illuminated by 561 nm and 405 nm lasers. The intensity of the 405 nm activation laser was adjusted by using continuous density filter (Thorlabs). For tracking, traces were obtained using the TrackMate plugin (33) for Fiji. The tracking was performed using simple LAP tracker with max linking distance of 0.5 μm. Gap-closing max distance was set to 0.5 μm and gap-closing max frame gap was chosen to be 2. Further, the data was analysed using TrackR package in R as described in (23).

In DNA-PAINT experiments the 561 nm and 645 nm lasers were sequentially switched while acquiring 20,000 frames in each channel at 80 ms exposure time. The molecule localizations were obtained using ThunderSTORM plugin (48) in Fiji. Then the localization files were additionally filtered in SMAP (49) where groups of localizations featuring less than 10 events were discarded and the localizations located only within the 100 nm z-slice in the middle part of bacteria were kept. Further analysis was performed on a z-projection of the filtered data.

### FLIM-FRET

Time-correlated single-photon counting FLIM measurements were performed on a home-made two-photon excitation scanning microscope based on an Olympus IX70 inverted microscope with an Olympus 60× 1.2NA water immersion objective operating in de-scanned fluorescence collection mode. Ti:Sapphire oscillator (Mai Tai DeepSee, Spectra Physics - 80 MHz repetition rate, ≈ 70 fs pulse width) at 10-20 mW was used for two-photon excitation at 930 nm. Emitted photons were collected through a 680 nm short pass filter (F75-680, AHF, Germany) and a 525/50 nm band-pass filter (F37-516, AHF, Germany) and directed to a fibre-coupled avalanche photo-diode (SPCM-AQR-14-FC, Perkin Elmer) connected to a time-correlated single photon counting (TCSPC) module (SPC830, Becker & Hickl, Germany). More details on the FLIM FRET setup are given in references (27, 50). The data analysis was performed using a commercial software (SPCImage, V8.1, Becker & Hickl, Germany).

## Acknowledgements

HM’s PhD grant was supported by a governmental fellowship. Y.M. is grateful to the ‘Institut Universitaire de France (IUF)’ for its support and the additional research time provided. We acknowledge the Imaging Center PIQ-QuESt (https://piq.unistra.fr/), a member of the national infrastructure France-Bioimaging supported by the French Research Agency (ANR-10-INBS-04).

## Supplementary Note 1: Table S1

**Table 2.**
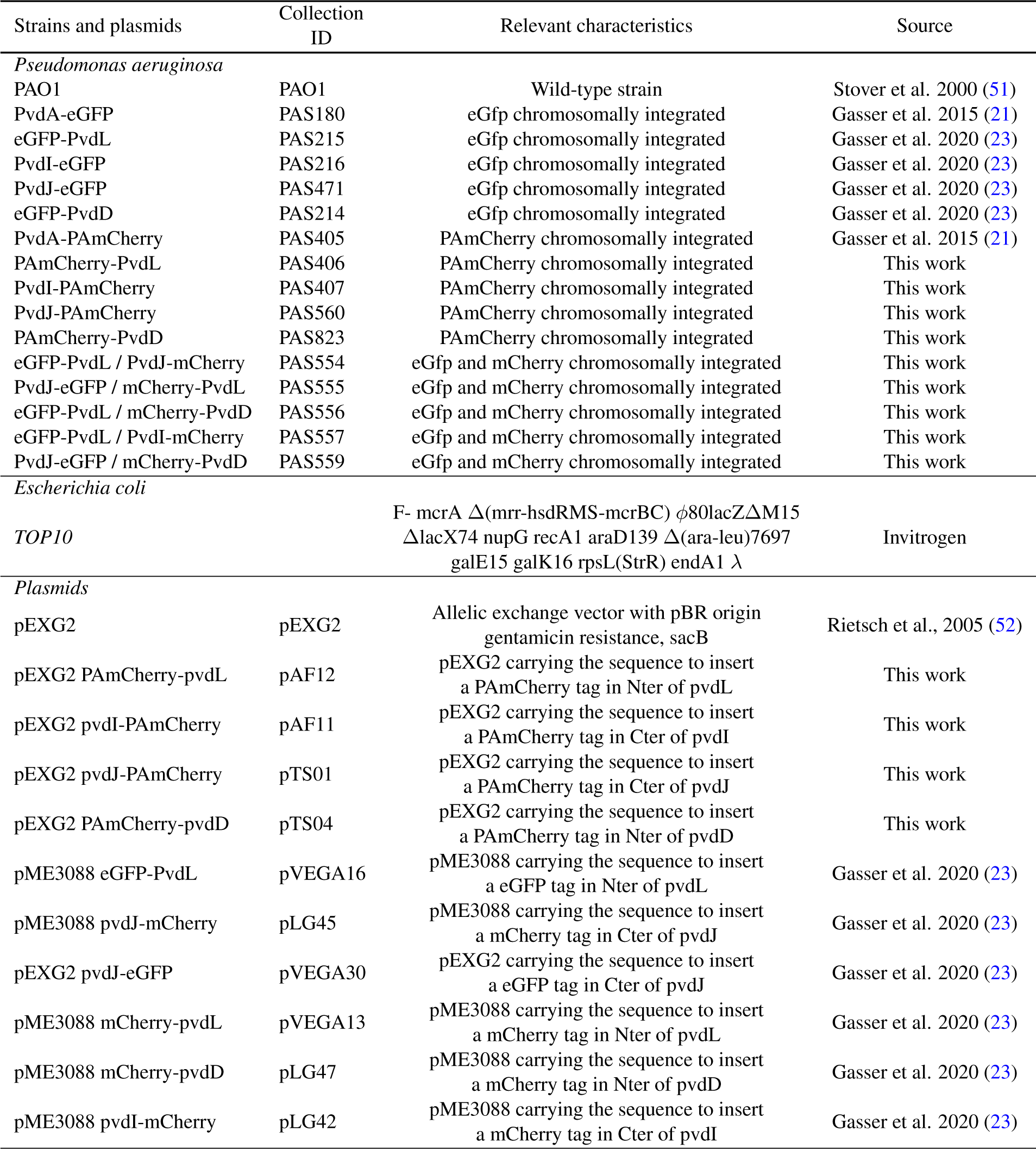
Strains and plasmids used in this work.

## Supplementary Note 2: Table S2

**Table 3.**
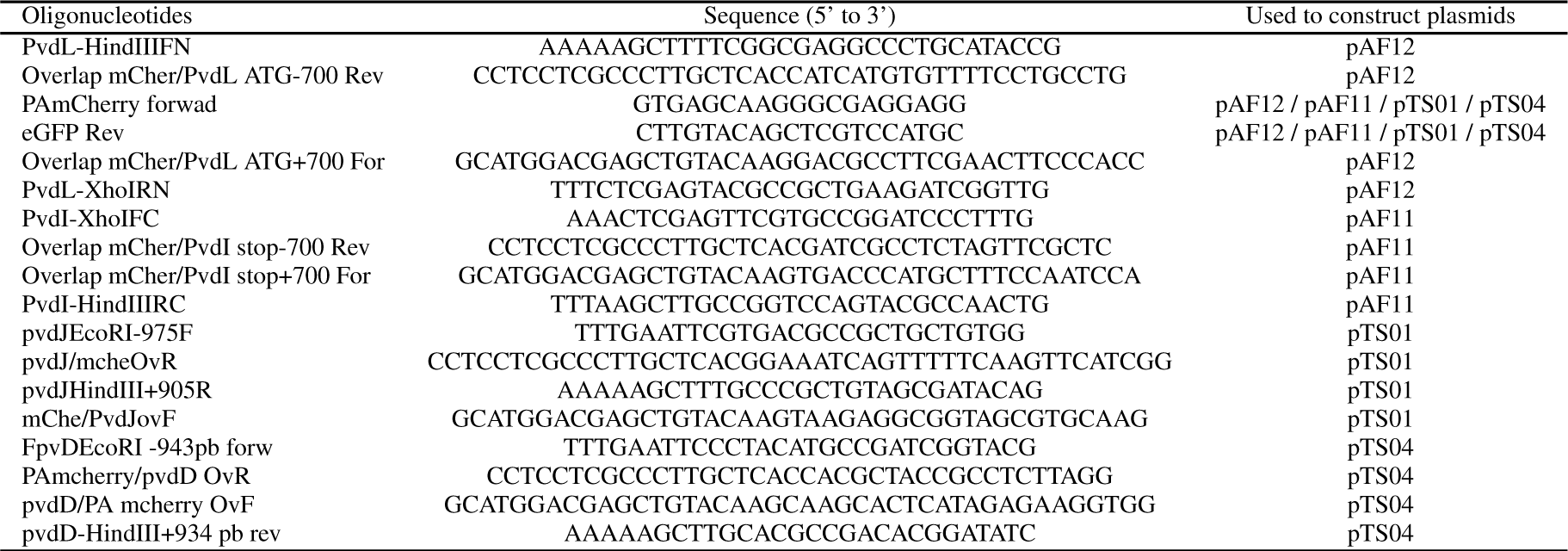
Oligonucleotides used to construct plasmids.

## Supplementary Note 3: Figure S1

**Fig. 6.**
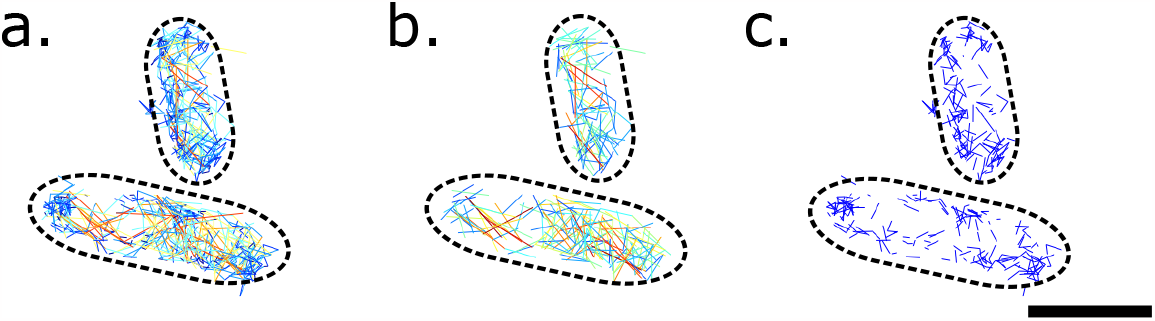
Representative diffusion maps of PvdD (the scale bar = 1 *μ*m) made of **a**. all the reconstructed trajectories, **b**. trajectories filtered by a median jump distance *>* 0.2 *μ*m or **c**. trajectories with median jump distances shorter than 0.15 *μ*m.

